# Decontaminating genomic data for accurate species delineation and hybrid detection in the Lasius ant genus

**DOI:** 10.1101/2024.11.27.625433

**Authors:** Kristine Jecha, Guillaume Lavanchy, Tanja Schwander

## Abstract

Advancements in genetic technologies have allowed us to generate large data sets relatively quickly and easily. However, without proper quality control checks, the inferences drawn from such data can be erroneous and go on to misinform further studies. DNA contamination between focal samples of the same or closely related species can have major impacts on downstream analyses, but their presence is seldom tested. Here, we created a pipeline combining competitive mapping to remove reads from intergeneric contamination, followed by a filtering method using allelic depth ratio frequencies to exclude intrageneric contamination. We then used a RADseq dataset of over 1,000 Swiss *Lasius* ants that were cross contaminated to various levels prior to sequencing to assess the impact of contamination on inferences of introgression. The original dataset presented widespread introgression between species in which hybridization has never been recorded. After thorough decontamination, we found only one individual with a strong signature of introgression, between the species *L. emarginatus* and *L. platythorax,* revealing that introgression is extremely rare in this genus. Implementing our method of filtering can significantly improve the robustness of biological findings based on genomic datasets. We recommend that systematically checking for the presence of cross contamination should be a key step in the preprocessing of genomic datasets.

## Introduction

The rise in use of DNA sequencing technology in recent years has boosted research in taxonomy and evolution of non-model species. Notably, it enabled researchers to validate and complement morphology-based species descriptions (Schlick-Steiner et al., 2010), identify cryptic species (Seifert et al., 2017), and uncover cases of hybridization and introgression (Taylor and Larson, 2019). However, an important aspect of this technology that should not be ignored is the risk of cross-contamination, i.e., contamination stemming from different samples used within the same research project, and the downstream consequences that it can cause if not properly addressed. Notably, cross-sample contaminations can mimic gene flow, which can result in overestimating hybridization and introgression between species.

While the issue of contamination in genome assemblies, typically involving microbial or dietary DNA of the sequenced organism (Eisenhofer et al., 2019; Merchant et al., 2014; Zhou et al., 2024), is now well recognized (Breitwieser et al., 2019; Boothby et al., 2015; Goudey et al., 2022; Koutsovoulos et al., 2016; Laurin-Lemay et al., 2012), similar awareness has not yet permeated studies relying on population-level genomic data. As for genome sequencing, contamination between samples may be introduced prior to sequencing, such as during the sample collection, in-laboratory sample preparation, or during the sequencing process itself (Ballenghien et al., 2017; Prüfer, & Meyer, 2015; Qi et al., 2021).

While contamination between distant taxa can be relatively easy to identify and remove (Cornet & Baurain, 2022; Laetsch & Blaxter, 2017; Schmieder, & Edwards, 2011), the more closely related the sample of interest and the contaminant are, the more difficult it can be to detect contamination. This poses a problem for population genomic datasets, which are based on many samples of the same or related species, and where cross-contamination can be particularly problematic. Thus, genetic delineation of species and studies of introgression between species can be easily obfuscated if cross-contamination is present in the data set. The ability to detect contamination and clean datasets for downstream analyses can be key, especially in cases where the initial sampling effort was very large or resampling would be difficult to impossible.

Here, we develop a two-step pipeline to decontaminate population genomic data. The first step leverages competitive mapping to eliminate reads originating from distant species. The second step is based on allelic depth ratio (ADR) and detects and eliminates contaminations from conspecifics and closely related species. We apply it to a dataset of over 1,000 *Lasius* ants from the canton of Vaud in Switzerland. These samples were part of a larger citizen science initiative aimed at surveying ants of several genera of the region using RADseq for genotyping in combination with COI gene analysis (Avril et al., 2019; Freitag et al., 2020; Szewczyk et al., 2024; Lavanchy et al. 2025). However, the RADseq data set of *Lasius* samples appeared to have widespread contamination that needed to be purged in order to salvage any information from this data. *Lasius* species have been notoriously difficult to distinguish with many cases of cryptic species that are morphologically similar and cases of misclassification later leading to unjustifiably synonymizing (Seifert, 1991; Seifert 1992; Wilson, 1955). Additionally, while there have been several reports of hybridization between *Lasius* species (Feldhaar et al., 2008; Pearson, 1983; Seifert, 1992; Seifert, 2019; Van der Have et al., 2011), a more encompassing investigation into the frequency of hybridization and introgression between the various species within the genus has never been performed.

## Methods

### Study area and sampling

We collected samples of *Lasius* as part of the *Opération Fourmis* citizen sampling campaign (Avril et al., 2019; Freitag et al., 2020) as well as through a structured sampling process throughout the Vaud canton of Switzerland in 2019 (Szewczyk et al., 2024). About 10 workers from each colony were collected and stored in 70% EtOH until DNA extraction. In total 3,173 *Lasius* worker ants were collected, 434 from the structured sampling, and 2,739 from citizen sampling (Fig. 1). These samples were identified morphologically by expert taxonomists to the species level when possible following Seifert (2018).

**Figure 1.**
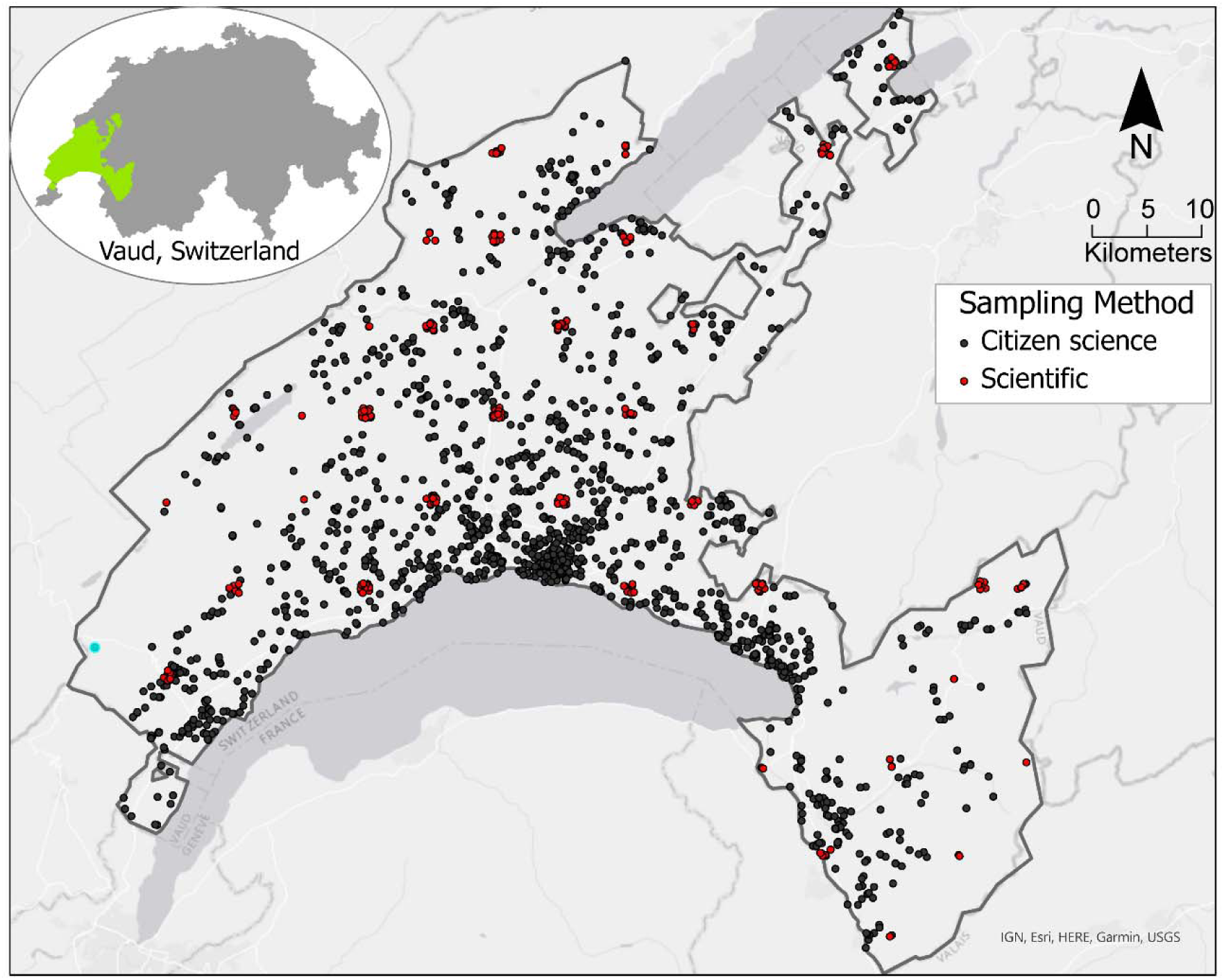
Map of *Lasius* colony sample locations within the canton of Vaud in Switzerland (n =3,138). Black points represent ants collected by the citizen science *Opération Fourmis* project, red points represent ants collected via structured scientific sampling. 35 samples missing geographical data were not included in the map.

### DNA extraction and sequencing

For each colony, we selected one worker ant at random to undergo genetic sequencing. For each worker, three legs were used for DNA extraction. Using only legs allowed us to preserve the remainder of the ant as a voucher specimen. We genotyped up to 200 individuals for each (morphologically identified) species. For species that had more than 200 colonies collected in the field, we down-sampled those from the citizen science campaign that were near each other to ensure geographically widespread sampling, resulting in a total of 1,171 genotyped individuals. The DNA extraction and COI barcoding procedures were outsourced to the Canadian Center for DNA Barcoding (CCDB) (Guelph, Canada) for 790 samples and to AllGenetics (A Coruña, Spain) for the remaining 381. A fragment of the COI gene was successfully amplified using primers LepF1 and LepR1 for 1,111 samples (Hebert et al., 2004). The specific protocols used for extraction, PCR amplification, and sequencing are available at the CCDB’s website https://ccdb.ca/resources (last checked Nov 17, 2023). We generated nuclear DNA sequences using RAD sequencing protocols described in Brelsford et al. (2016). We used restriction enzymes EcoRI and Msel to digest the genomic DNA followed by ligating custom barcode adaptors to the DNA fragments, which were then amplified in 16 PCR cycles. Fragments with a size of 280 to 430 bp were selected using 2% agarose cassettes on a BluePippin (Sage Science). We sequenced these libraries on eight Illumina HiSeq 2500 lanes (single end, 150 bases) to generate the final read data.

### SNP calling

We demultiplexed the sequenced reads using stacks v2.53 process_radtags command with the options -c -q -r -t 143 (Catchen et al., 2013) and mapped the reads against an available reference genome of *Lasius niger* (BioProject: PRJNA1159026; Masson et al., 2024) with the mem algorithm of BWA v0.7.17 (Li & Durbin, 2010). Alignment file formatting and summaries were created using samtools v1.15.1 (Danecek et al., 2021). We built loci using gstacks with options --phasing-dont-prune-hets and --ignore-pe-reads and ran populations with the option --write-single-snp to retain only one SNP per RAD locus. We were then able to filter the data with vcftools 0.1.14 (Danecek et al., 2011) keeping only genotype calls with a minimum depth of 8 and loci with a minor allele count of 2 which were present in at least 75% of individuals. Any individuals missing more than 50% of the genotyped loci were excluded. As a quality control, we looked at the relationship between the proportion of missing genotypes and “relative heterozygosity”, measured as the proportion of heterozygous alleles to total genotyped SNPs.

### Competitive mapping

In order to decontaminate the RADseq reads, we used a competitive mapping approach inspired by Feuerborn et al. (2020): concatenating the reference genomes of species representing potential contaminants and the target reference genome, mapping the RADseq reads to the concatenated genomes, and removing reads mapping to the non-target genomes (Figure 2A, B). In our case, we combined the *Lasius niger* genome with ant genomes from seven genera to represent other ant genera also found in Vaud, Switzerland as part of the *Opération Fourmis* sampling (Szewczyk et al., 2024). These included *Camponotus*, *Formica*, *Leptothorax*, *Myrmica*, *Tapinoma*, *Temnothorax*, and *Tetramorium* (Romiguier et al., 2022 and BioProjects: PRJNA445978, PRJNA819254, and PRJNA533534). We mapped the reads from each sequenced *Lasius* individual to this multi-genus concatenated genome with the mem algorithm of BWA v0.7.17 (Li & Durbin, 2010). We could then separate the reads into those preferentially mapping to the target *Lasius* genome and those that were mapping to a possible contaminant genome using R v4.2.1 (R Core Team, 2021) by identifying the species’ genome that every read in each sample mapped to in their sequence alignment file and retaining a list of the each read name that mapped either primarily or secondarily to the *Lasius* genome. To filter out possible contaminants, we used the “filterbyname.sh” function of BBMap v38.63 (Bushnell, 2014) with this read list as the “name” input and the “include=t” parameter. We then remapped these selected reads to the *Lasius niger* reference genome alone and performed SNP calling as previously stated.

**Figure 2.**
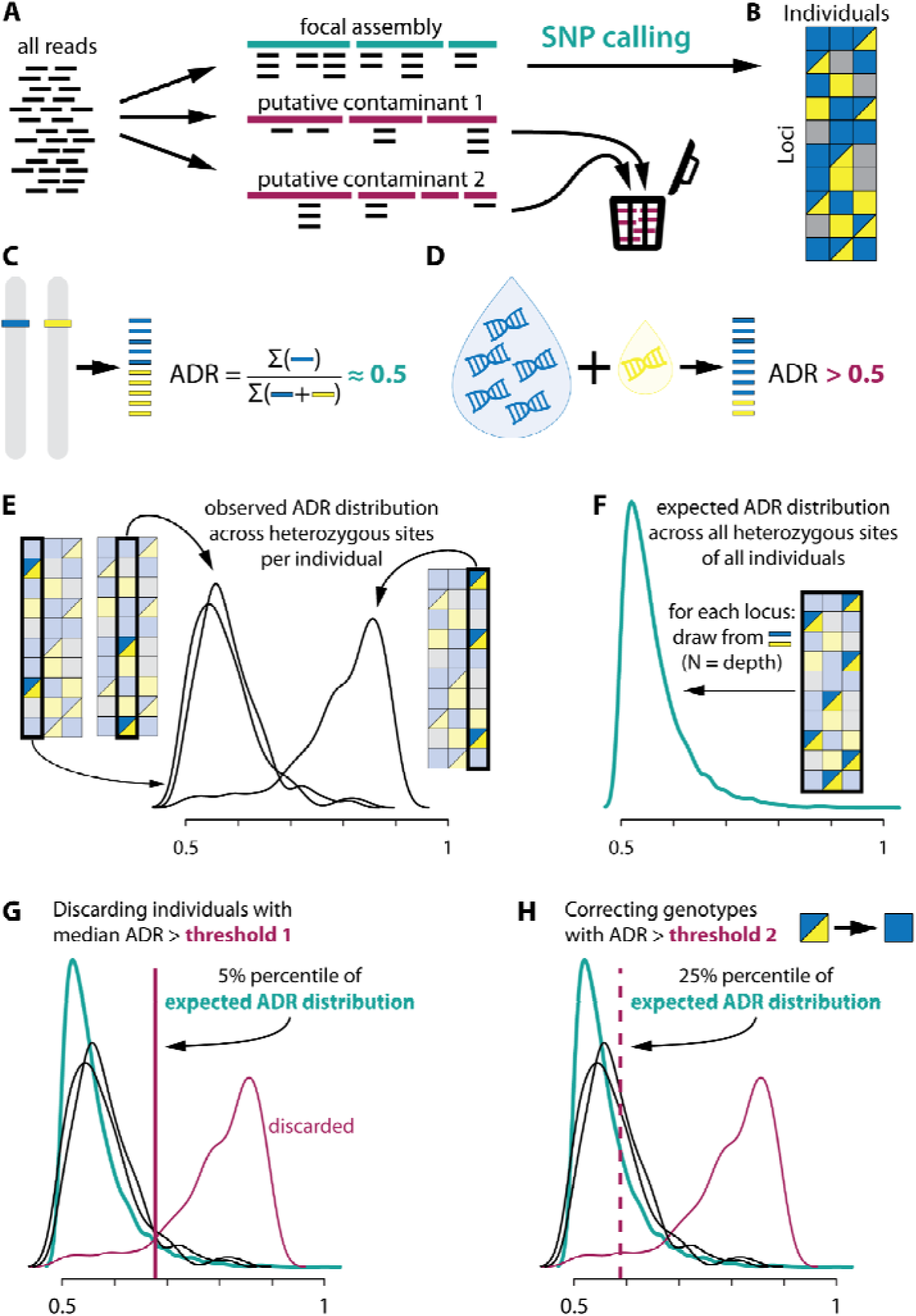
Overview of the decontamination pipeline. **A** Competitive Mapping. Reads are mapped simultaneously to the focal genome assembly and assemblies of putative contaminant taxa. **B** Typical dataset after SNP calling. Blue and yellow are two alleles, grey is missing data. **C**-**G** Allelic Depth Ratio filtering approach. **C** Allelic Depth Ratio (ADR) is expected to be around 0.5 for a real heterozygote. **D** Contaminations are expected to lead to ADR higher than 0.5. **E** We first estimate ADR distributions across all heterozygous loci for each individual independently. **F** We then estimate the expected ADR distributions across all heterozygous loci given observed depths. **G** We discard individuals whose median ADR is beyond the top 5% percentile of the expected ADR distribution. **H** We correct genotypes with ADR beyond the top 25% percentile of the expected ADR distribution by removing the allele with the fewest reads.

### Allelic depth ratio filtering

Putative contamination between samples from the same genus cannot be removed by our competitive mapping approach. We therefore used an additional, complementary approach based on allelic depth ratio. We define allelic depth ratio (ADR) at a heterozygous locus as the ratio of reads of the allele with most reads to the sum of all reads at the locus. It can thus range from 0.5 to <1. For a real heterozygote in a diploid individual, it is expected to be around 0.5 in all single-copy loci (Fig. 2C). However, if a heterozygous genotype is caused by contamination, the amount of contaminant DNA is expected to be lower than the original sample’s DNA, and thus ADR will be higher than 0.5 (Fig. 2D). We used ADR-based filtering in two steps. First, we discarded individuals with a highly skewed ADR distribution, as these most likely represent highly contaminated samples. We estimated the expected distribution of ADR by sampling from a binomial distribution with locus depth as the number of observations for all heterozygous sites in all individuals. We then considered that individuals whose observed median ADR (across all their heterozygous loci) was beyond the top 5% percentile of this expected ADR distribution were highly contaminated and were discarded (Fig. 2G).

Second, in the retained individuals we corrected the individual genotypes with ADR beyond the top 25% percentile of the expected ADR distribution. We assumed that the allele with the fewest reads was derived from a contaminant and considered the individual homozygous for the other allele (Fig. 2H). This filtering was performed using a custom script in R v4.2.1 (R Core Team, 2021) with the package vcfR v1.14.0 (Knaus & Grünwald, 2017).

### Haploid allele filtering

As an alternative way to remove contaminants and assess the overall robustness of our findings, we artificially “haploidized” our competitively mapped dataset. At all heterozygous sites, we artificially removed the allele with the fewest reads. When there were the same number of reads at both alleles, one was drawn at random. We also considered homozygous sites haploid. This filtering was performed using a custom script in R v4.2.1 (R Core Team, 2021), using the package vcfR v1.14.0 (Knaus & Grünwald, 2017).

### Molecular-based species identification

To investigate the congruence between the morphological identification and SNP-based identification, we used the SNP calls to create a MultiDimensional Scaling (MDS) plot with the R package MASS (Venables & Ripley, 2002) with R v4.2.1 (R Core Team, 2021). In conjunction with the MDS clustering, we ran ADMIXTURE (Alexander et al., 2009) in 10 replicates for K values ranging from 1 to 15. We then selected the best value of K and the best replicate based on Cross-Validation error (K=9). We conducted the same analysis in haploid mode with the option --haploid=“*” for the “haploidized” dataset. We then visualized the results with pong (Behr et al., 2016) and chose the most representative run. In cases where the morphological identification was ambiguous or different from the genetic identification, the individual was reassigned to its genetic species identification. We were able to directly identify hybrid individuals using the ADMIXTURE data results. Any individual that contained 98.4375% or less of their main species ancestry (expected ancestry proportion after 5 generations of backcrossing) was identified as being of hybrid ancestry.

With the COI gene sequences, we created a phylogeny of the *Lasius* individuals. We first aligned the sequences using the L-INS-i algorithm of mafft v7.481 (Katoh et al., 2005). We then built a Maximum Likelihood tree using IQ-TREE v2.2.0.5 (Minh et al., 2020) using ModelFinder Plus (Kalyaanamoorthy et al., 2017) to select the best substitution model, and assessed it with 1000 UltraFast Bootstraps. Species delineation from the COI gene was compared to combined RADseq and morphological delineations to further confirm species identity for each individual.

## Results

### Decontamination of RADseq Data

RADseq genotyping was performed on 1,171 *Lasius* individuals. After excluding individuals missing at least 50% of genotypes, 981 individuals were left. The allelic depth ratio showed signs of a large amount of nuclear DNA contamination as many individuals exhibited an ADR peak around 0.9 (Fig. 3A). The effect of contamination can also be seen by the dramatically increased proportion of heterozygous loci, especially in individuals that are missing a higher proportion of genotypes (Fig. 3B). In order to obtain biologically meaningful results based on the SNP data created from these individuals, it was necessary to decontaminate the data.

**Figure 3.**
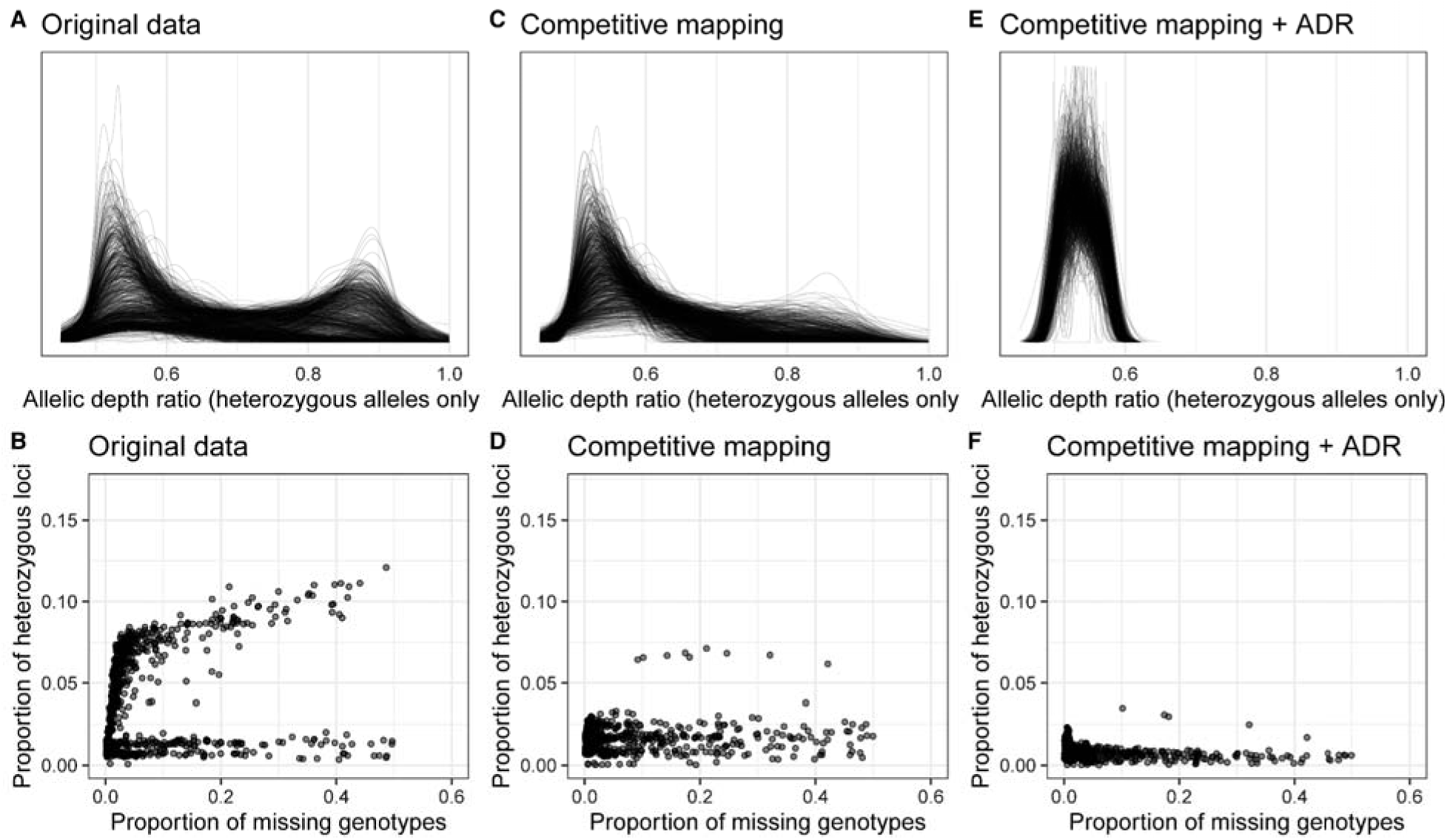
**A** Allelic depth ratio distribution of contaminated RADseq data. **B** Proportion of heterozygous loci per individual in the contaminated RADseq data. **C** Allelic depth ratio distribution of the competitively mapped RADseq data. **D** Proportion of heterozygous loci per individual in the competitively mapped RADseq data. **E** Allelic depth ratio distribution in the filtered RADseq data (filtered based on the allelic depth ratio, ADR). **F** Proportion of heterozygous alleles per individual in the ADR-filtered RADseq data.

Through competitive mapping, it appeared that the majority of the read contamination originated from ants from the *Formica* genus as the majority of non-*Lasius* reads mapped to the *Formica* genome (supplemental Fig. S1). After filtering out the reads that did not preferentially map to the *Lasius* genome, an average of 64.3% (median 72.6%, standard deviation 18.1%) of reads remained. After SNP calling and discarding low-data individuals, we retained 974 samples.

The efficiency of competitive mapping as a decontamination technique can be seen clearly when comparing the decontaminated allelic depth ratio (Fig. 3C) and the decontaminated proportion of heterozygous loci per individual (Fig. 3D) to the contaminated data (Fig. 3A,3B).

However, competitive mapping is only able to remove contaminants outside of the *Lasius* genus, and intrageneric contamination would remain. To eliminate this type contamination, we compared a technique of artificial “haploidization” of the samples and an allelic depth ratio (ADR) filtering. For the latter, we discarded individuals considerably outside the expected ADR and corrected individual genotypes that were slightly beyond the expected distribution (Fig. 3E and 3F). After SNP calling and removing low-data individuals, the ADR filtering retained 902 individuals.

In order to identify which filtering technique was successful at eliminating the most contamination while still being useful for molecular analysis, we compared allelic depth ratio plots of each filtering technique (Fig. 3). Competitive mapping alone was not sufficient to filter all contaminants, since some individuals still showed signatures of contamination (ADR peaks >> 0.5). The competitive mapping in conjunction with either ADR filtering or “haploidization” were the two best options for removing contaminants based on allelic depth frequency and MDS clustering (Fig. 4 B-E). However, the “haploidization” resulted in more loss of information and would preclude additional analyses such as estimations of heterozygosity. For this reason, we selected the competitive mapping + ADR filtering method for downstream analyses.

**Figure 4.**
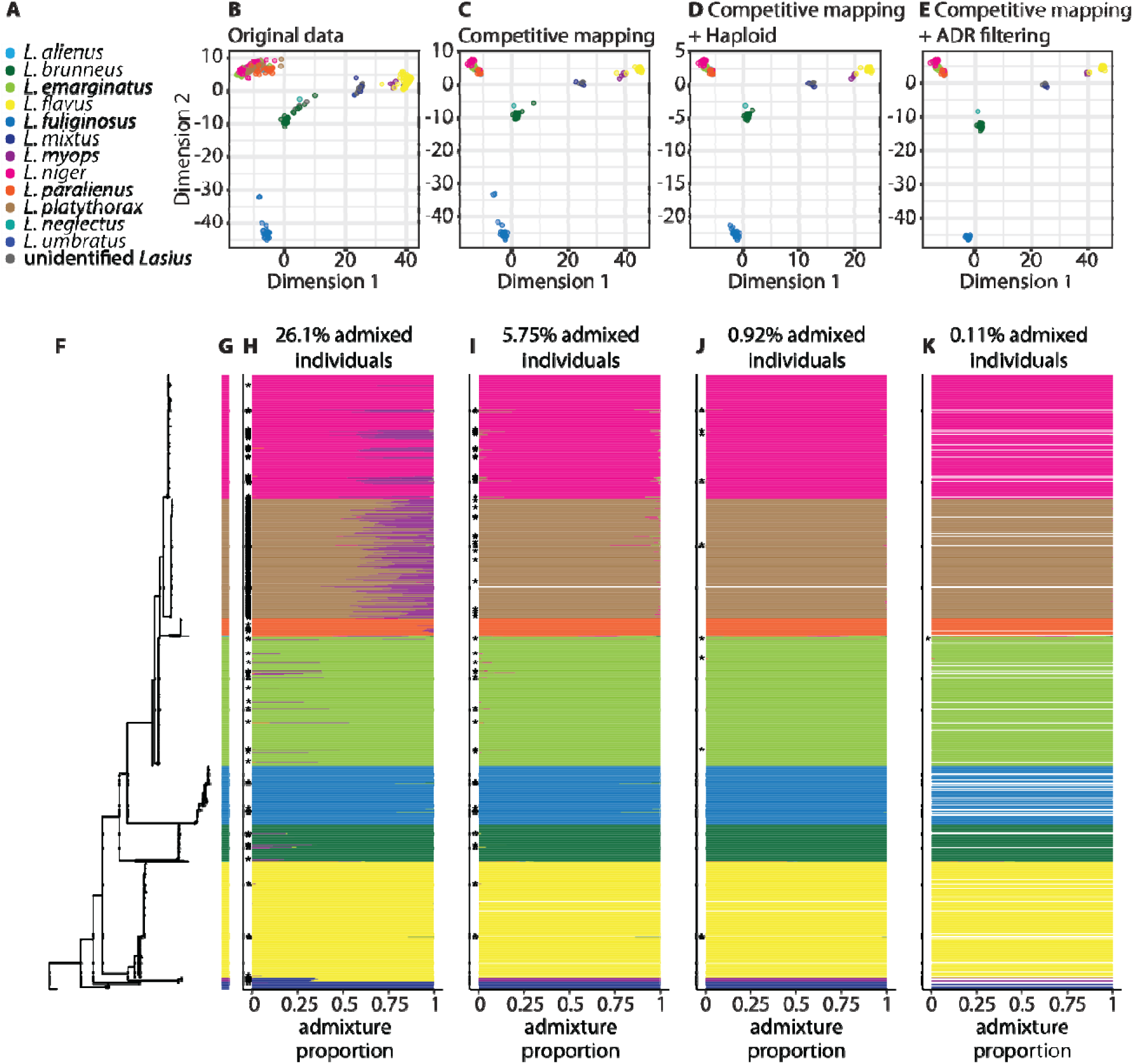
Taxonomic inference based on the different versions of the dataset. **A** Species color codes. **B** - **E** MultiDimensional Scaling (MDS) plots of original data and each decontamination process. **F** Phylogeny of the COI mitochondrial gene. **G** Final genetic-based species identifications. **H - K** ADMIXTURE proportions of original data and each decontamination process, corresponding to the adjacent MDS plots. Stars denote individuals inferred to be introgressed. White lines correspond to individuals that were filtered out due to high levels of suspected contamination or low remaining SNP data. **B & H** Original (contaminated) data. **C & I** Competitive mapping only. **D & J** Competitive mapping + haploidization. **E & K** Competitive mapping + Allelic Depth Ratio filtering.

In an effort to identify a possible source of the contaminations, we examined the percentage of contaminated samples processed by the two sequencing centers. Thus, we counted all samples that were removed from the original dataset due to insufficient coverage after the removal of contaminant reads or if they had a highly skewed ADR that would indicate high contamination. While such highly contaminated samples were found from both sequencing centers, contaminated samples originated from CCDB nearly twice as often than from AllGenetics, see Table 1.

**Table 1.** Outcome of samples from each company used for DNA extraction in the final competitively mapped + ADR filtered dataset.

| Sequencing center | Number of samples | Percentage of samples |
| --- | --- | --- |
| **AllGenetics** | 355 total |  |
| kept | 337 | 94.9% |
| removed | 18 | 5.07% |
| **CCDB** | 626 total |  |
| kept | 565 | 90.3% |
| removed | 61 | 9.74% |

### Species Delineation

Out of the 902 genotyped individuals that had sufficient data after competitive mapping and ADR filtering, 22 could not be confidently identified to the species level during initial morphological identification. The availability of nuclear genotype information in conjunction with COI sequence information allowed us to unambiguously assign these samples to a species. Additionally, 18 morphological misidentifications could be corrected based on the molecular analyses; see Supplementary Table 1 for misidentification reassignments.

Three individuals, two morphologically identified as *L. alienus* and one *L. neglectus*, could not confidently be assigned to a genetic group in the ADMIXTURE analysis (Fig. 4E) due to the low number of conspecifics. For the two *L. alienus,* the ancestry proportions are unique enough that it is unlikely that they belong to any of the larger species’ groups. For the sole *L. neglectus* individual, in the absence of other samples of this species, it remains unclear if this is truly a *L. neglectus* or another *Lasius* species not represented in the remaining dataset. But its phylogenetic position as a sister group to *L. brunneus* is consistent with the position of *L. neglectus* in the *Lasius* phylogeny from Steiner et al. (2004).

The combination of the MDS, COI, and ADMIXTURE analyses also allowed us to investigate the presence of cryptic species. MDS plots based on SNP distances were created for each species with at least 10 individuals to improve the resolution for detecting cryptic species (Supplementary Fig. 2). However, there was no evidence of genetic subdivision within any of the species in the MDS plot, COI phylogeny, nor when the K parameter was increased in the ADMIXTURE analysis, indicating that there are no cryptic species within the sampled *Lasius* species.

MDS, ADMIXTURE, and COI species delineations were always consistent except in one case where an individual morphologically identified as *L. platythorax* clustered with other *L. platythorax* in the MDS and had a majority of *L. platythorax* SNPs according to the ADMIXTURE analysis; however, this individual was on the *L. emarginatus* branch of the COI phylogeny (Fig. 4A). The ancestry proportion of this individual is 95.06% from *L. platythorax* and 4.94% from *L. emarginatus* based on ADMIXTURE. This individual appears to have retained the mitochondria of a female *L. emarginatus* during a hybridization event in its ancestry, followed by generations of backcrossing into the *L. platythorax* population, resulting in a predominantly *L. platythorax* nuclear DNA.

### Hybrid identification

All individuals with admixture percentages of at least 1.56% (the expected proportion for a hybridization event followed by five generations of backcrossing) were classified as “admixed” (hereafter referred to as “hybrids”; Fig. 4C-E). Before filtering, 256 individuals were identified as hybrids using this cut-off. However, read filtering by competitive mapping reduced the number of hybrids to 56 individuals, artificial haploidization of the data in conjunction with competitive mapping reduced this to 9 individuals, and allelic depth filtering with competitive mapping resulted in only 1 individual classified as admixed with our threshold. The hybrid ancestry of this individual is further supported by its mismatch of nuclear and mitochondrial DNA. It exhibits a high percentage of *L. platythorax* nuclear DNA, but its COI genotype clusters with *L. emarginatus* samples. This difference between the number of admixed individuals after the haploidization method and ADR method of filtering is because false hybrids identified in the haploidized dataset were removed during the ADR filtering due to falling above the 5% percentile of expected ADR distribution. The admixed identification of these individuals in the haploidized dataset indicates that at certain loci, more than half of the reads at that site contained an allele originating from contamination. However, as the overall allelic depth ratio frequency of these individuals was high enough for them to be excluded during the ADR filtering, this is likely a result of contamination at sites with low coverage. Additional individuals also showed signs of admixture after decontamination, but with ancestry of the donor species lower than 1.56%, suggesting older introgression events.

## Discussion

### Sources and consequences of contamination

Contamination can arise at multiple steps of wet-lab processing and has substantial potential to mislead downstream analyses. External DNA may be introduced through gut content, parasites, microbiota, or accidental contact with other samples. It can inflate genome size estimates or create spurious signatures of horizontal gene transfer (Boothby et al., 2015; Crisp et al., 2025). Cross-contamination among samples of the same or closely related species poses additional challenges: it can mimic introgression, inflate estimates of connectivity and heterozygosity, or even generate artificial paralog-like signals if more than two haplotypes appear in a presumed diploid individual. These risks increase sharply when input DNA is limited. In our case, DNA was extracted from only three legs of a single ant. This is perhaps an extreme situation, but necessary in species-delineation studies where voucher specimens must remain intact for morphological examination.

Despite these risks, contamination screening remains rare in population-genomic workflows. While genome assembly pipelines commonly incorporate such checks (e.g., BlobTools; Laetsch & Blaxter, 2017), similar precautions are seldom applied to RADseq or resequencing datasets. To address this gap, we developed a two-step procedure combining competitive mapping with allelic depth ratio (ADR) filtering. This approach enabled us to detect and remove contamination from both congeneric and conspecific sources, and from more distantly related ant taxa.

### Identifying contaminations

Distinguishing genuine biological patterns from contamination is challenging, and overly stringent contamination filtering risks inadvertently removing true signal. We therefore examined alternative explanations for the high numbers of hybrids and skewed ADR distributions in our initial dataset.

PCR bias could in principle distort ADRs in hybrids, but such an effect would require consistent bias across homologous loci in related species while not affecting within-species polymorphisms. We consider this implausible. PCR bias could in theory be caused by major differences in GC content between related species; however, GC content is nearly identical across the eight publicly available *Lasius* genomes (36.43–36.89%; Vizueta et al., 2025).

RADseq-specific biases such as differential restriction enzyme activity due to epigenetic modifications could also theoretically produce skewed ADRs. However, this would require persistent, genome-wide epigenetic differences between hybridizing species maintained across multiple generations of backcrossing, which is highly improbable.

Species-specific paralogs represent another potential source of misleading ADRs. Yet for recently evolved paralogs to bias genome-wide patterns, they would need to occupy a substantial portion of the genome, generating large genome-size differences among closely related species. This is inconsistent with known patterns in ants, where genome size variation is mostly attributable to repeats (Vizueta et al., 2025), which are typically underrepresented in RADseq because repetitive regions are removed during size selection and SNPs in such regions are filtered due to high coverage. Collectively, these considerations support contamination as the most parsimonious explanation.

### Data decontamination

Our pipeline effectively removed contamination from the RADseq dataset, enabling robust genetic characterization of over 900 *Lasius* individuals. Competitive mapping efficiently eliminated reads originating from non-target ant genera, while ADR filtering removed residual interspecific contamination within *Lasius*. Although this procedure resulted in the exclusion of some low-coverage samples, it produced a substantially cleaner dataset that more accurately reflects the genetic structure of *Lasius* populations in the canton of Vaud, Switzerland.

Competitive mapping is particularly powerful for removing exogenous DNA, but its effectiveness depends on the availability of high-quality reference genomes for both focal taxa and potential contaminants. We used *L. niger* as the representative *Lasius* genome, the most complete assembly available, which could theoretically bias retention toward closely related species. However, we found no evidence of such bias in this genus (Supplementary Fig. S2). As more reference genomes become available, the power and precision of this approach will continue to increase.

ADR filtering is broadly applicable to any type of genomic dataset on any organism and is not restricted to RADseq, but thresholds should be adjusted in systems where species differ substantially in paralog content. Duplications followed by divergence can produce ADR patterns resembling contamination (e.g., AAAB genotypes with ADR = 0.75), particularly in low-coverage samples. Users should therefore tailor cutoffs to the genomic architecture of their study organisms.

Interestingly, most contamination in our dataset appeared to originate from samples processed at one of the two sequencing centers involved in DNA extraction. This mirrors earlier reports of contamination issues linked to certain providers (e.g., Ballenghien et al., 2017). This highlights the need for rigorous quality control, even when working with established sequencing centers, and for routine contamination checks.

### Insights into Lasius taxonomy and patterns of introgression

After decontamination, one individual with an admixture proportion of >1.56% (indicative of five generations of backcrossing) was detected among the 902 individuals: an advanced-generation backcross between *L. platythorax* and *L. emarginatus*. This individual exhibited ∼95% *L. platythorax* nuclear ancestry but carried *L. emarginatus* mitochondria, consistent with an initial cross between a female *L. emarginatus* and a male *L. platythorax*, followed by several generations of backcrossing. This case highlights both the potential for rare but fertile hybridization and the importance of combining nuclear and mitochondrial data; COI alone would have misclassified this individual.

Although previous work suggested frequent hybridization in *Lasius* (Seifert, 1992, 2019; Feldhaar et al., 2008), our broad but unbiased sampling revealed no F1 hybrids, suggesting that hybridization may not be as common as previously believed.

We also used genetic data to validate or correct morphological identifications. Nuclear genotypes and COI sequences revealed no cryptic lineages in species with enough individuals (at least 10) to investigate it (*L. brunneus, L. emarginatus, L. flavus, L. fuliginosus, L. niger, L. paralienus,* and *L. platythorax*). This contrasts with earlier periods of taxonomic revision in *Lasius* (Seifert, 1991; Seifert, 1992; Wilson, 1955) and illustrates the value of integrating morphological and molecular approaches (Dayrat, 2005).

For species represented by few samples, assignments remain less certain. Nonetheless, the placement of *L. alienus* near *L. paralienus* and *L. neglectus* near *L. brunneus* in the COI tree is consistent with past taxonomic work (Seifert, 1992; Steiner et al., 2004). *L. neglectus* was revealed to be present in the Vaud region for the first time (Cremer et al., 2008; Espadaler et al., 2007), an important finding for monitoring this invasive species. Salvaging this contaminated dataset thus yielded biologically meaningful insights that would otherwise have been lost.

Contamination can convincingly mimic introgression, mask species boundaries, inflate heterozygosity, or produce patterns resembling three-way hybridization. Our preliminary analysis of contaminated *Lasius* samples initially suggested extensive intergeneric introgression with *Formica* and hybridization among multiple *Lasius* species, patterns that, while possible, would be extraordinary in nature and inconsistent with known reproductive barriers. While similar patterns have been reported in other taxa, genetic inference alone risks being misinformed due to undetected contamination (Moore & Coulson, 2020; Lancaster et al., 2006; Rothfels et al., 2015; Thörn et al., 2024; Toews et al., 2018; Zbinden et al., 2023). Explicit testing for contamination is therefore essential before interpreting extraordinary hybridization signals.

The method presented here is broadly applicable to any genetic data, including RADseq, resequencing, and other genotyping methods. While it cannot substitute for careful wet-lab practices, it provides a rigorous safeguard against misleading conclusions. Incorporating contamination screening into standard genomic pipelines will improve the accuracy, reliability, and interpretability of population-genomic data across biological systems.

## Data Availability

The code used and scripts created for the data analysis can be found on GitHub: https://github.com/glavanc1/Lasius_species_ID. Demultiplexed RADseq reads can be found on SRA (Bioproject PRJNA1176511). COI sequences can be found on GenBank (accession 6036-7160). See Supplementary Table 3 for SRA and GenBank accession numbers for each Lasius individual and outcome during the contamination filtering process.

## Acknowledgments

We thank Marc Bastardot, Marjorie Labédan, Christine La Mendola, Heather Ryder and Simon Vogel for their help processing samples. This study was supported by funding from a bequest of M. Rullens to the University of Lausanne, the fondation Herbette, the Société Vaudoise des Sciences Naturelles, and the Retraites Populaires.

