## Supplementary figures and images for "Decontaminating genomic data for accurate species delineation and hybrid detection in the Lasius ant genus"

### Supplementary Figure S6

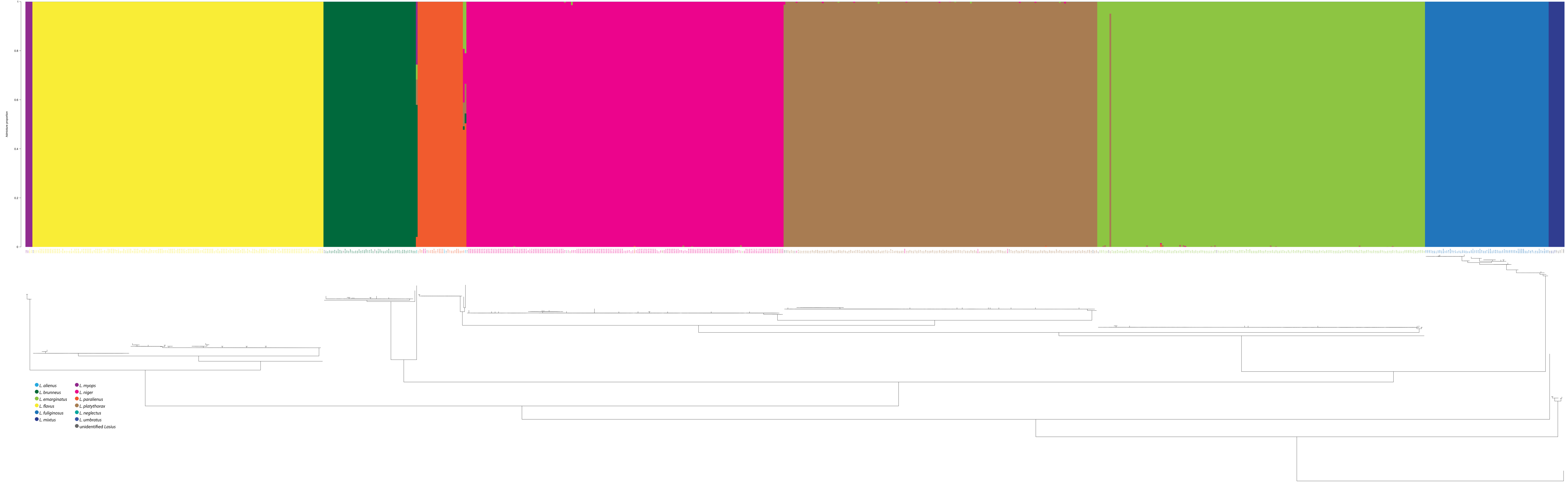
