## Supplementary Figures and Tables for "Decontaminating genomic data for accurate species delineation and hybrid detection in the Lasius ant genus"

### Supplementary Material

#### Supplementary Table 1. Individuals with incongruent morphological and genetic species identification. Species reassigned based on genetic species

| **ID** | **Morphological Species** | **Genetic Species** |
| --- | --- | --- |
| 9990311 | *L.alienus* | *L.paralienus* |
| 7451 | *L.emarginatus* | *L.niger* |
| 16418 | *L.flavus* | *L.myops* |
| 9993114 | *L.flavus* | *L.myops* |
| 1960 | *L.niger* | *L.platythorax* |
| 9990390 | *L.niger* | *L.platythorax* |
| 9990699 | *L.niger* | *L.paralienus* |
| 9990779 | *L.niger* | *L.paralienus* |
| 9991831 | *L.niger* | *L.platythorax* |
| 9991944 | *L.niger* | *L.platythorax* |
| 12831 | *L.paralienus* | *L.alienus* |
| 9990347 | *L.paralienus* | *L.niger* |
| 9990517 | *L.paralienus* | *L.niger* |
| 10548 | *L.platythorax* | *L.niger* |
| 18867 | *L.platythorax* | *L.niger* |
| 5957 | *L.platythorax* | *L.niger* |
| 12012B | *L.umbratus* | *L.mixtus* |
| 9990311 | *L.alienus* | *L.paralienus* |

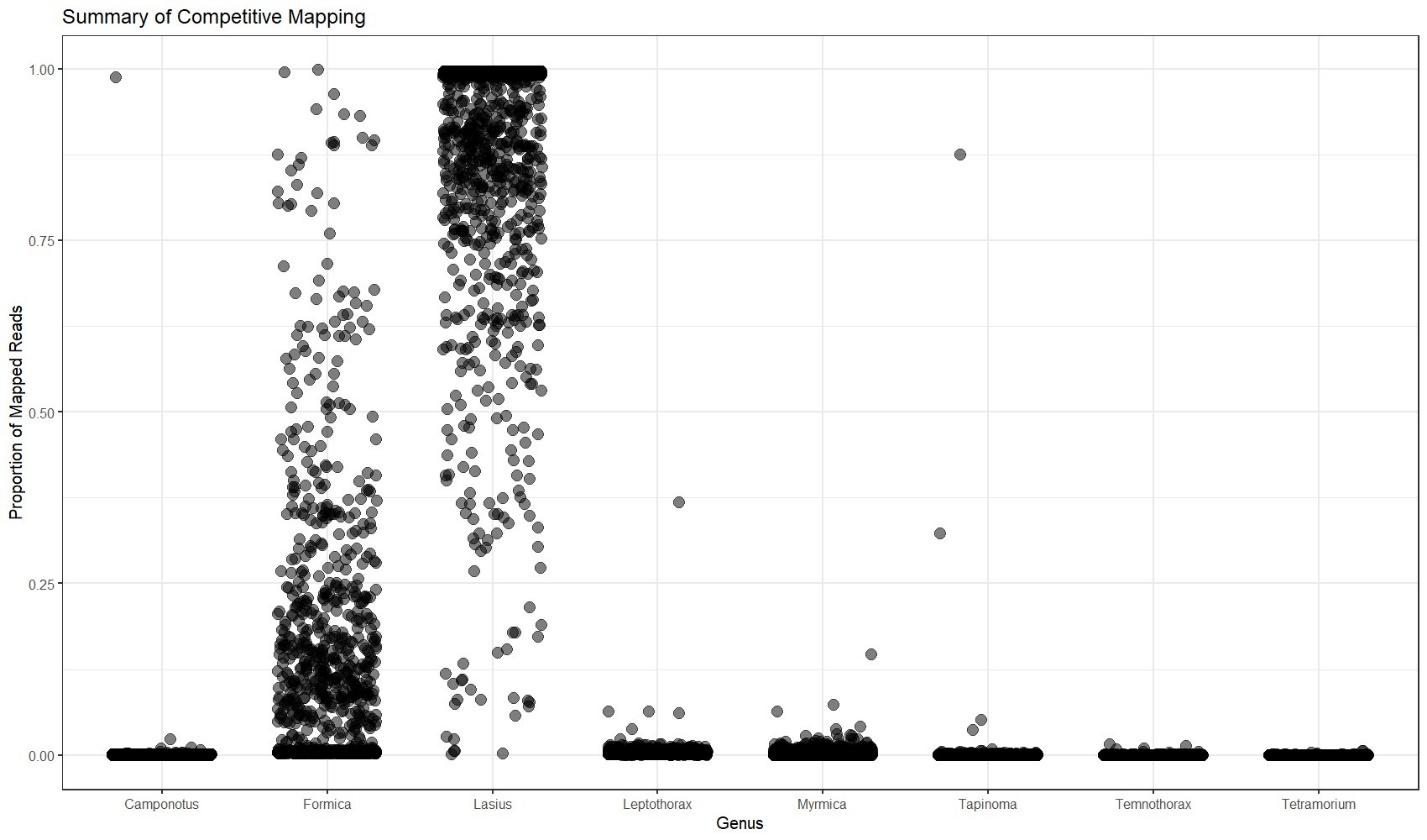

Supplementary Figure 1. Results of competitive mapping *Lasius*

RADseq reads from *Lasius* individuals competitively mapped against a concatenated genome of eight ant species from different genera. Proportion of reads from each sample that mapped to each of the genomes shown. Apparent contamination of reads from *Formica* species or closely related genus.

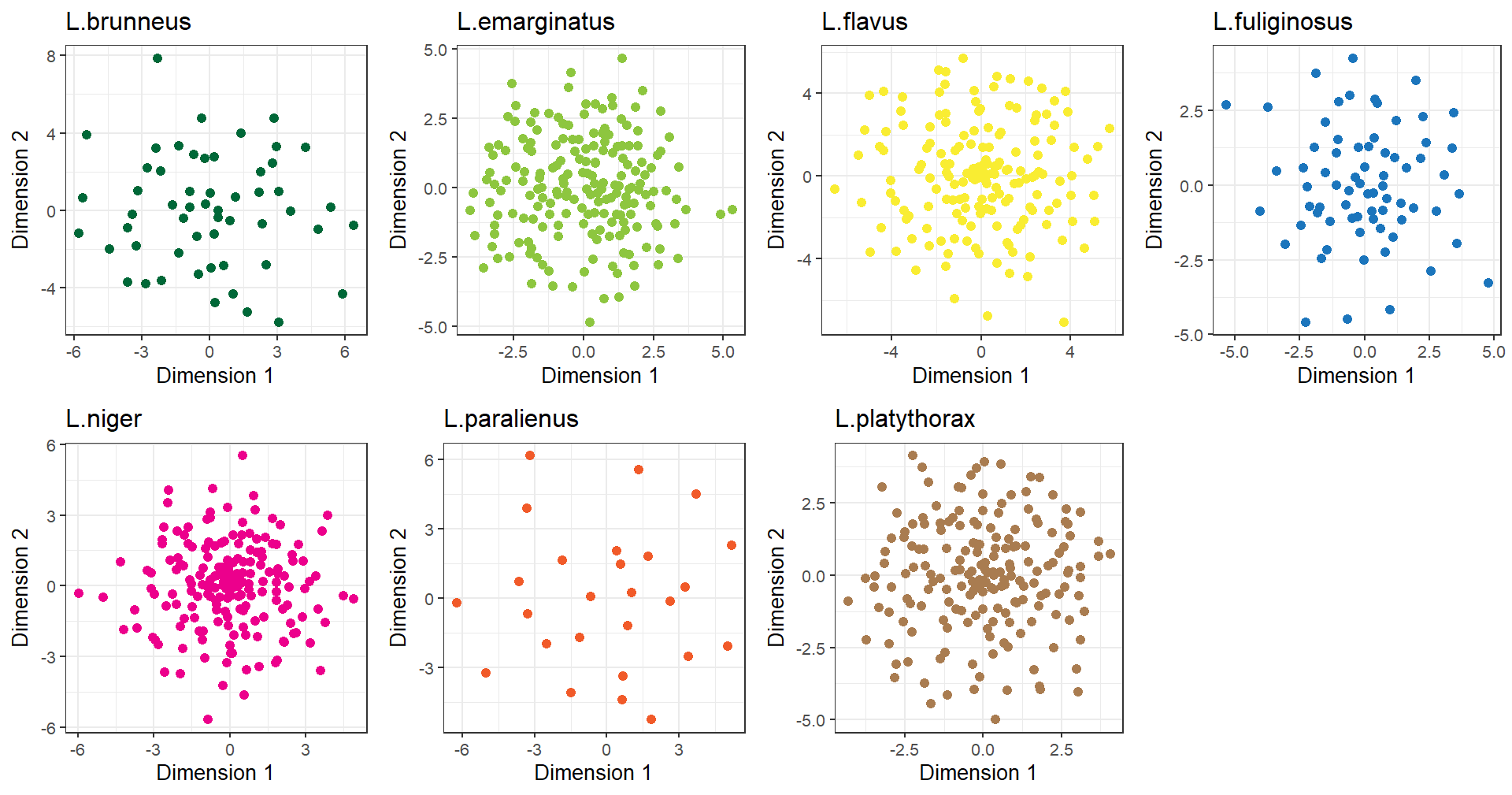

Supplementary Figure 2. Verification of no cryptic species within *Lasius* species

MDS plots for each *Lasius* species with more than ten individuals to investigate the presence of cryptic species. No sub clustering within the species indicates no apparent cryptic species.

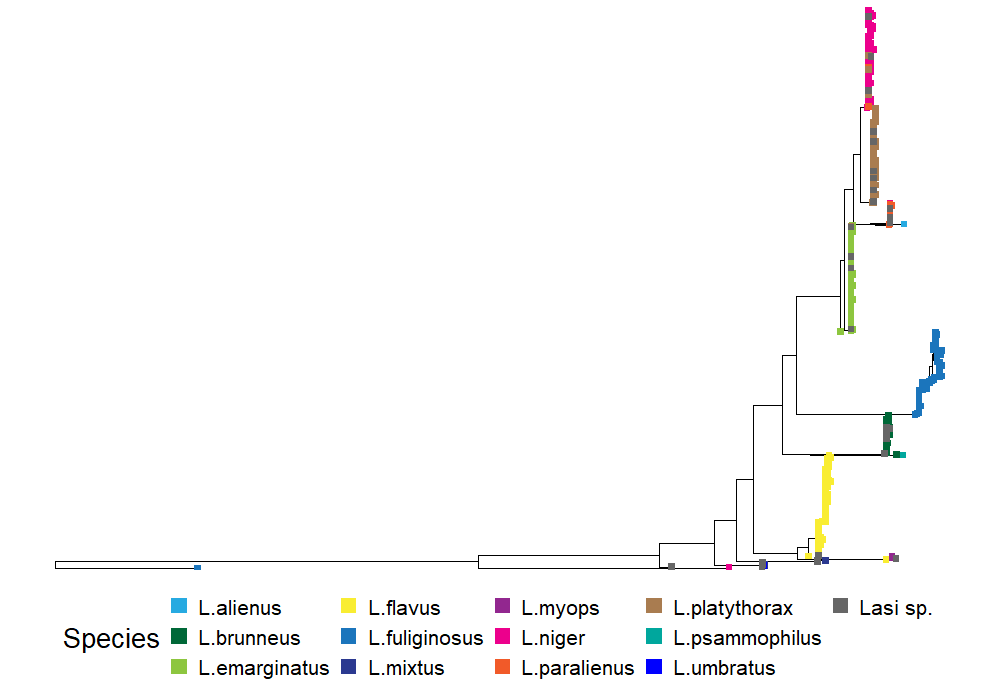

Supplementary Figure 3. Complete phylogeny of COI gene among *Lasius* individuals. Tip colors based on original morphological identification (n=1,111).

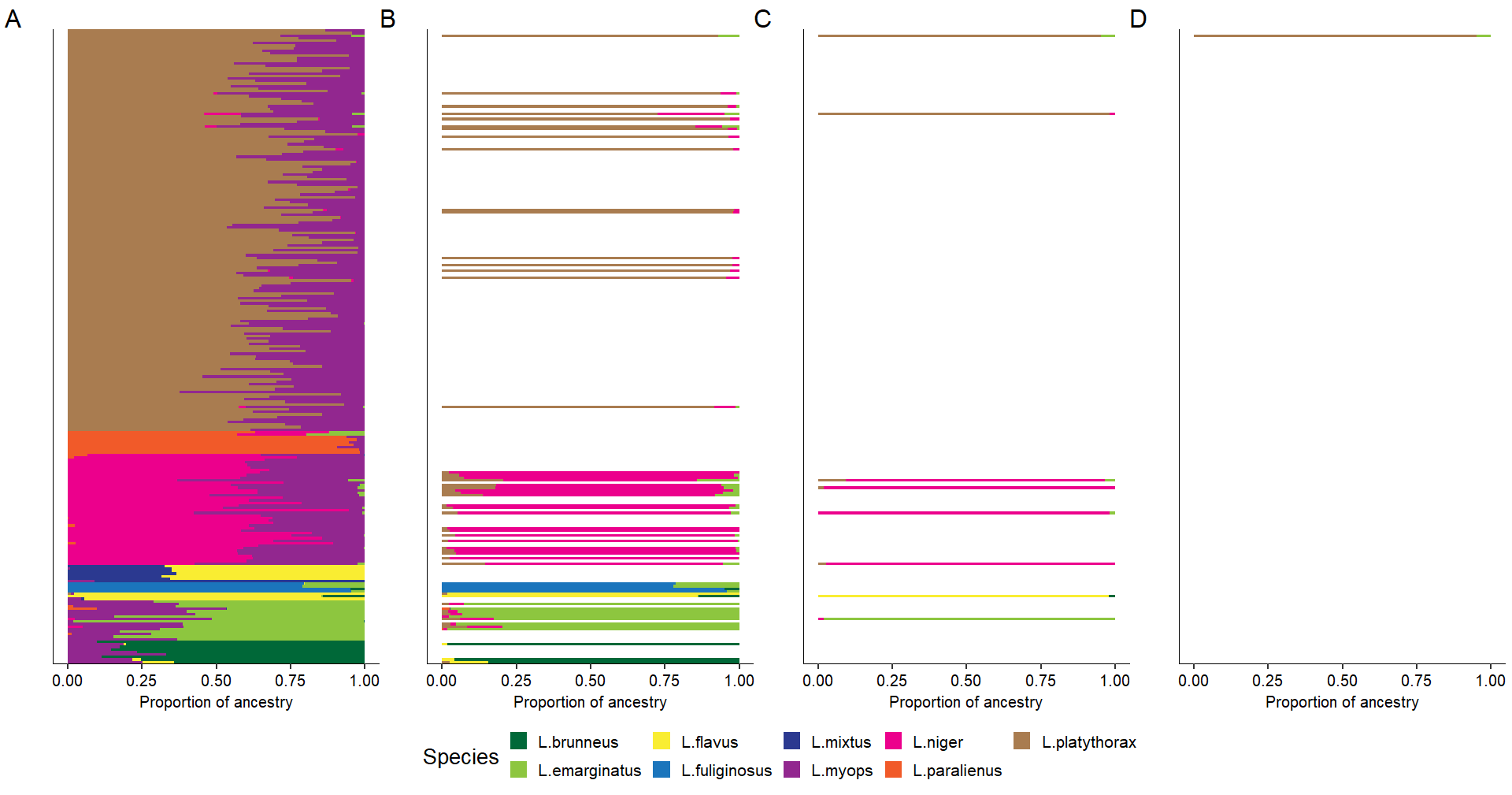
Supplementary Figure 4. ADMIXTURE plot of admixed individuals A) ADMIXTURE plot of potential hybrids from the original data (n=260). Each horizontal line represents the proportion of ancestry from one individual aligned to the COI phylogeny. B) ADMIXTURE plot of potential hybrids from data after competitive mapping (n=54). C) ADMIXTURE plot of potential hybrids from data after competitive mapping and artificial “haploidization” (n=7). D) ADMIXTURE plot of potential hybrids from data after competitive mapping and ADR filtering (n=1).

Supplementary Figure 5 (uploaded separately). Admixture plot (top), morphological identification (colored label) and COI phylogeny (bottom) for all 902 individuals for which both nuclear and COI data were available. Admixture barplot, sample name and phylogeny tips are aligned. Colors as in Figure 4.
